# Intercellular adhesion molecule-1 reprograms microglia to improve cognitive functions by inhibiting ERK/STAT3 signalling pathway in a model of Alzheimer’s disease

**DOI:** 10.1101/2024.05.20.594921

**Authors:** Soumita Goswami, Subhas Chandra Biswas

## Abstract

**Background:** Microgliosis is one of the early hallmarks of Alzheimer’s disease (AD) that plays a crucial role in AD-associated neuroinflammation. However, strategies to harness microglial activation without affecting its homeostatic function are currently lacking. Our recent finding revealed astrocyte secreted cytokine Intercellular adhesion molecule 1 (ICAM-1) improves memory and cognitive impairment in 5xFAD mice model of AD. Here, we investigated the possible mechanism of ICAM-1 that involves modification of microglial activation state and function.

**Methods:** We treated primary microglia cultures with amyloid-ß (Aß) in presence or absence of ICAM-1 and then assessed microglial inflammatory activation and phagocytic clearance of Aß by western blotting and confocal imaging. We further treated 5xFAD mice with ICAM-1 intraperitoneally and ICAM-1 antagonist lifitegrast intranasally to assess the beneficial effect of ICAM-1 on Aß mediated microgliosis, synaptic protein expressions and cognitive deficits.

**Results:** Here we report that, ICAM-1 inhibits microglial inflammatory activation by inhibiting ERK-STAT3 pathway which is indispensable for microglial inflammation. Moreover, ICAM-1 was found to potentiate microglia to engulf and eliminate Aß in primary culture and reduced Aß plaque load and associated microglial activation in 5xFAD mice hippocampus. This reduced plaque levels and associated microgliosis in turn refurbished synaptic protein expressions and improved cognition and memory in these mice. Interestingly, ICAM-1 mediated microglial modification along with cognitive improvement was partially lost when ICAM-1-LFA-1 interaction was inhibited.

**Conclusion:** Taken together these findings delineate the importance of ERK/STAT3 pathway in Aß mediated microglial inflammation and the modulatory role of ICAM-1 in microglial activation and phagocytosis in order to improve clearance of Aß and associated memory and cognitive impairments.

## Background

Amyloid-β (Aβ) deposition, being the prime event in Alzheimer’s disease (AD) development, induces neuroinflammation which manifested majorly by aberrant microgliosis along with induction of associated pro-inflammatory proteins, that play substantial role in degeneration of neurons and its synapses leading to cognition and memory problems in AD patients (Hardy & Allsop, 1991; Heppner et al., 2015; Perlmutter, 2016). Genome wide association studies (GWAS) have identified several inflammation associated genes such as CLU, CR1, TREM2, CD33 majorly expressed by microglia as risk genes for AD strongly suggesting a close association between microglia mediated neuroinflammation and AD (Bradshaw et al., 2013; Guerreiro et al., 2013; Jonsson et al., 2013; Lambert et al., 2009). In healthy brain microglia detects any pathological abnormalities like misfolded proteins, dying neurons, protein aggregates by binding to pathogen associated molecular pattern (PAMP)s or damage associated molecular pattern (DAMP)s with the help of different pattern recognition receptors (PRR) present on its surface. Binding of Aβ oligomers and fibrils to scavenger receptors like CD68, CD36-CD47-α61β integrin complex, TLR-4, TLR-6 activates microglia to phagocytose Aβ and produce pro-inflammatory cytokines and chemokines as an innate immune response (Bamberger et al., 2003; Stewart et al., 2010). Aβ plaque deposition results when Aβ synthesis overweighs its clearance. In sporadic AD, improper Aβ clearance due to the challenged phagocytosis induces the excessive production of inflammatory mediators that create a very detrimental condition for neurons leading to disease development (Hickman et al., 2008). Conventional anti-inflammatory drugs such as cyclooxygenase inhibiting non-steroidal anti-inflammatory drug (NSAID)s were found to inhibit inflammation associated AD pathogenesis at early stages but were found to interfere with the phagocytic clearance of Aβ in advanced AD conditions. Immunomodulatory therapies must target microglial inflammation while improving their ability of phagocytic clearance simultaneously maintaining their homeostatic functions. Therefore, it is important to properly identify and define the regulators of homeostatic and inflammatory microglia responses.

Early activation of glial cells often engages different mitogen activated protein kinase (MAPK) pathways as upstream signalling regulators (Cargnello & Roux, 2011). In mammals, three major MAPKs are identified namely Extracellular regulatory kinase (ERK), c-Jun NH2 terminal kinase (JNK) or stress related protein kinase (SAPK) and P38 (Kaminska, 2005; Plotnikov et al., 2011). Our lab previously shown the essential involvement of p38 and JNK MAPK activation in Aß mediated astrogliosis (Saha et al., 2020). Moreover, ERK and its downstream transcription factor, signal transducer and activator of transcription 3 (STAT3) activation has been found to be involved in glial activation and subsequent inflammation (Chen et al., 2021; Huiliang et al., 2021). Upon binding of growth factor ligands to receptor tyrosine kinases, ERK gets phosphorylated and activated which further phosphorylates STAT3 at serine 727 residue. In macrophage cells STAT3 phosphorylation specifically at serine 727 was found to be involved in LPS induced production of inflammatory cytokines like TNF-α and IL-1β (Balic et al., 2020). Thus, this pathway could act as an important target for therapy in controlling the aberrant microgliosis in AD.

Recently we and others have shown the beneficial effect of astrocyte secreted Intercellular adhesion molecule 1 (ICAM-1) in reduction of Aβ levels by inducing neprilysin production and in improving memory and cognitive deficits in AD models (Guha et al., 2022; Kim et al., 2012). ICAM-1, a transmembrane glycoprotein belonging to immunoglobulin super family is also secreted by microglia cells in different neurological disorders (Chu et al., 2023; Lee & Benveniste, 1999; Müller, 2019). ICAM-1 majorly functions in leukocyte migrations during inflammation in a variety of diseases by binding to its cognate receptor Lymphocyte function associated antigen 1 (LFA-1) with its 190 aa long LFA-1 binding region in the I domain (Gorina et al., 2014; Wee et al., 2009). Additionally, ICAM-1 has been found to induce phagocytosis in LPS primed macrophages (Zhong et al., 2021).

In this study, we specifically investigated the effect of ICAM-1-LFA-1 interaction in modifying microglia induced neuroinflammation and associated neurodegeneration in a transgenic AD mouse (5xFAD). We identified ERK-STAT3 pathway as a novel and indispensable mechanism that underlies microglial inflammation in presence of Aβ. Moreover, our data first time revealed the immunomodulatory effect of ICAM-1 on Aβ mediated microgliosis leading to improvement in synaptic health and cognitive impairments.

## Materials and Methods

### Materials

Aβ1-42 was purchased from Alexotech (Vasterbotten County, Sweden), 1,1,1,3,3,3 hexafluoro-2-propanol (HFIP), Dimethyl sulfoxide (DMSO), U0126, Bovine serum albumin (BSA), Trypsin, Insulin, poly-D-lysine, glucose, ammonium per sulphate (APS), β-mercaptoethanol (β-ME), RIPA buffer, TEMED, citric acid, paraformaldehyde (PFA), Twin 20, Triton X 100, were from Sigma-Aldrich (St. Louis, USA). Dulbecco’s modified Eagles medium (DMEM), Fetal Bovine Serum (FBS), goat serum, Penstrep, prolong gold antifade with DAPI were purchased from Thermo Fisher Scientific (MA, USA). Recombinant ICAM-1, Lifitegrast, ICAM-1 ELISA kit and cytokine array kit were from R&D systems. Cell culture dishes, plates, filter units, sterile pipettes, flasks, tubes were purchased from BD Falcon (Schaffhausen, Switzerland) and Corning (NY, USA). PVDF membrane was purchased from GE Healthcare (Buckinghamshire, UK). Enhanced chemiluminescence (ECL) substrate, protein ladder, stripping buffer were purchased from Takara Bio (Kusatsu, Japan). Sodium dodecyl sulphate (SDS) was purchased from Merck (Darmstadt, Germany). Bovine Serum Albumin (BSA), TRIS buffer, glycine were purchased from Sisco Research Laboratories Pvt. Ltd (Mumbai, India). All other fine chemicals were procured from standard local suppliers unless stated otherwise. List of antibodies used for this work, their dilutions for each type of application and sources are given in the Additional file 1.

## Methods

### Preparation of Amyloid-β (Aβ) oligomer

Lyophilised Aβ peptide was reconstituted in 100 % HFIP to a final concentration 0.1 μM and centrifuged in a speed vac (Eppendorf, Hamburg, Germany) to completely remove HFIP. The peptide pellet was carefully dissolved in 5mM anhydrous DMSO with the help of a Hamilton syringe and the solution is sonicated in 37°C water bath for 10 mins. The peptide solution is aliquoted and kept in −80°C for future use. Just before use, oligomerisation is performed by treating with 0.1% SDS and 1x PBS in 37°C for 16-18 hours followed by dilution with 1x PBS to get a final concentration of 100 μM and incubated for 18-24 h at 37°C. The Aβoligomers are run in SDS-PAGE to check accuracy in process.

### Mixed glia culture

Mixed glial cultures were prepared from Sprague Dawley rat pups aged 1-2 days following the protocol of (Garwood et al., 2011; Saha & Biswas, 2015). Briefly, equal numbers of male and female rat pups were taken and after decapitation brains were isolated and kept in 1x of Hanks balanced salt solution (HBSS) buffer. After successfully isolating the cortices, they were roughly chopped and incubated with a solution of trypsin and BSA in 37°C for 20 mins. The brain pieces were triturated with pasture pipette and passed through a nylon mesh to prepare single cell solution. After centrifugation at 1200 rpm for 10 mins the pellet was dissolved in DMEM with10% FBS media and seeded at a density of 1.5 million per T25 flask and 0.75 million per well of 6 well plates. Mixed glia cultures were maintained in DMEM with 10 % FBS for 20 days while replenishing the medium every alternate day.

### Primary Microglia Culture

After maintaining the mixed glia culture in DMEM with 10 % FBS for 20 days microglia cells were isolated by mild trypsinisation according to the protocol mentioned by (Saura et al., 2003) with slight modifications. Mixed glial cells were incubated with trypsin-DMEM at a ratio of 1:8 at 37°C for 30 mins. After successful removal of other glial cells in the upper layer, the lower layer of pure microglia cells was maintained in glial conditioned medium for at least 24 h before any treatment.

### Treatment on cells

Rat primary microglia cells were treated with 1.5 uM Aβ oligomer with or without 100ng/ml of rrICAM-1 (R&D system, Minneapolis, USA) or anti-ICAM-1 antibody (Novus Biochemicals, USA) (4μl/ml) or MEK1/2 inhibitor U0126 (Sigma-Aldrich, St. Louis, USA) (10μM) and incubated for 1h, 3h, 6h, 16h and 24h.

### ELISA

Quantitative sandwich ELISA was performed using microglial conditioned medium before and after treatment with Aβ oligomer according to manufacturer’s instruction. Briefly, the ICAM-1 coated plates were incubated with microglial condition medium for 2h at room temperature in an orbital shaker. The plates were then washed with wash buffer and incubated with polyclonal anti ICAM-1 antibody conjugated with horseradish peroxidase (HRP) for 2h at room temperature on a shaker. The washing was repeated following addition of tetra methyl benzene and hydrogen peroxide mixture to the wells and kept in the dark at room temperature for 30 mins. Finally stop solution with hydrochloric acid was added and OD value was taken at 450 nm in a spectrophotometer and was plotted according to the standard.

### Cytokine array

Differential levels of 79 different cytokines were assessed using cytokine array kit (R&D systems) with primary microglia supernatant before and after Aβ exposure according to manufacturer’s instruction. Briefly, the antibody coated nitrocellulose membranes were incubated with blocking buffer for 1h at room temperature to block all the nonspecific binding on an orbital shaker. Samples were mixed with biotinylated detection antibody cocktail and the mixture was then added to the membranes and incubated over night at 4°C. On the next day the membranes were washed with wash buffer and incubated with the streptavidin horse radish peroxidase for 30 mins at room temperature on a shaker. After washing the membranes were incubated with a mixture of hydrogen peroxide and luminol to detect the proteins.

### Immunocytochemistry

After treatment, primary microglia cells were fixed with 4% para formaldehyde (PFA) in 1xPBS solution on glass coverslips for 10 mins. Cells were washed with 1xPBS and blocked with 3% normal goat serum for 1h at room temperature. Cells were immunolabelled with preferred primary antibody overnight at 4°C in a humid chamber. On the next day after washing with 1xPBS with 0.1% triton X-100 the cells were incubated with species specific secondary (alexa fluor) antibodies at room temperature for 1h. After washing, hoechst 33342 (Molecular Probes, Invitrogen, MA, USA) at a concentration of 1 μg/ml in 1x PBS solution was added for 30 mins at room temperature to stain the nuclei. The coverslips were mounted with Prolong gold antifade (Invitrogen, MA, USA) with DAPI and images were taken with Leica Sp8 STED confocal microscope (Wetzlar, Germany) at 63x magnification. Integrated density of staining, area of individual cell and the background fluorescence of individual image in different experimental conditions were derived using NIH-ImageJ software. The corrected total cell fluorescence (CTCF) was calculated as follows, CTCF = Integrated density – (area of selected cell x mean fluorescence of background readings).

### Animal housing and care

C57BL/6 and 5xFAD transgenic mice (both male and female purchased from Jackson Laboratory) with an average body weight of 25 - 30 g were kept separately in maximum three per individually ventilated cages (IVC) in a room with controlled temperature (24 ± 2°C), 12 to 12 h light-dark cycle, humidity (60 ± 5%) under the supervision of trained staff members in the animal house of CSIR-Indian Institute of Chemical Biology, Kolkata, India. All the studies were conducted in accordance with the guidelines formulated by the Committee for Control and Supervision of Experiments on Animals (CCSEA) of Department of Animal Husbandry and Dairying (DAHD), Ministry of Fisheries, Animal Husbandry and Dairying (MoFAH&D), Govt. of India with approval from the Institutional Animal Ethics Committee (IAEC).

### Treatment in animals

5xFAD and C57BL/6 mice of 5 months age were treated intranasally with lifitegrast (10 μM) and intraperitoneally with rrICAM-1 (1μg/kg body weight) with an interval of 1.5-2 h on every alternate day for 21 days. On 24^th^ day animals were sacrificed by CO2 inhalation and brains were collected in RIPA buffer (Thermo Fisher Scientific, West palm beach, USA). For immunohistochemical staining, the brains were collected after cardiac perfusion with 1xPBS to wash the blood channels and fixed with 4% PFA. The brains were kept in 4% PFA for 48 h and finally kept in cryoprotectant solution of 30% sucrose before cryo-sectioning. 20μm thin sections were prepared after freezing the brain at −29°C in the cryotome (Thermo Fisher Scientific, West Palm Beach, FL, USA) (Das et al. 2021). Brain sections were preserved in a cryoprotectant solution (30% ethylene glycol 20% glycerol in phosphate buffer Ph 7.4) at −20°C before immunostaining.

### Immunohistochemistry

20 μm thin brain sections were taken out of the freezer and washed 3-4 times with 1x PBS. Sections were subjected to permeabilization with 1xPBS with 0.4% triton X for 30 mins. After washing tissue sections were incubated with citric acid solution of pH 6 at 95°C for 10 mins for antigen retrieval. After washing, blocking buffer with 3% normal goat serum was added and incubated for 1h at room temperature. This was followed by immunolabelling with desired primary antibody diluted in blocking solution and kept 24-48h at 4°C on an orbital shaker. On the next day after washing, appropriate secondary antibody (alexa fluor 488/568) was added and incubated at room temperature for 1.5-2h. Lastly, the nuclei were stained with Hoechst 33342 (Molecular Probes, Invitrogen, MA, USA) solution at a concentration of 1 μg/ml in 1x PBS for 30 min at room temperature, followed by mounting with Prolong Gold Antifade with DAPI for microscopic analysis. Images were captured in Leica Sp8 STED confocal microscope (Wetzlar, Germany) at 63x magnification. Aβ plaque number and puncta number of synaptic proteins were calculated using puncta anlyzer plugin of NIH-imageJ software. The corrected total cell fluorescence (CTCF) was calculated by including integrated density, integrated density of background and area of the cell by ImageJ software. CTCF = Integrated density - (area of selected cell × mean fluorescence of background value).

### Behavioural Experiments

C57BL/6 and 5xFAD mice were randomly divided in four groups – WT (C57BL/6), 5xFAD, 5xFAD+rrICAM-1 (1μg/kg bodyweight) and 5xFAD+rrICAM-1+lifitegrast (10μM) and sequentially subjected to 3 behavioural tests naming open field test (OFT) or locomotion, novel object recognition (NOR) and cue dependent fear conditioning (CDFC). Animal numbers in each group are given in Additional file 4.

### Open Field Test (OFT)

OFT determines the general locomotory activity and exploratory behaviour in mice along with thigmotaxis. Thigmotaxis is an inherent tendency of animals to remain close to the walls as an act to reduce anxiety. Open-field test (OFT) apparatus (IR Actimeter, Panlab, Barcelona, Spain) consists of open arena made up of Plexiglas with photocell emitters and receptors positioned at equal distance along the perimeter of the arena (278×236×300 mm). During animal movement the infrared beam grid breaks and that can be recorded.

The animals were placed in an empty arena and allowed to explore for 10 mins, under dim lighting. The positions of the rodents were tracked using custom ActiTrack software (Panlab, Barcelona, Spain). The arena was divided into two zones; a square area occupying one-third of the total area in the center was considered as the ‘Inner zone’ and the rest the ‘Outer zone’. The total distance travelled (cm), the speed of the mouse (cm/sec) and the distance travelled in the outer zone (periphery) and that in the inner zone (center) were recorded. Distance travelled away from walls (time in center) were analyzed per 10-minute video to determine anxiety like behaviour.

### Novel object recognition (NOR)

It is a two-day experiment which majorly depicts hippocampal recognition memory function and explorative behaviour. During habituation mice were allowed to explore a rectangular box (50 cm x 30 cm) for 5 min on Day-1. On the same day, two identical (color and dimensions) objects were placed at parallel corners and the animals were kept to explore the objects for 10 mins. On probe day or day 2 one of the objects was replaced with an object of different color and shape and the mouse was returned to the box for 5 min. The time (sec) explored with novel and familiar objects on probe day were noted as Time with Novel (TN) and Time with familiar (TF) respectively and represented as exploration time (sec). Discrimination index (DI) and preference index (PI) were derived according to the formula – DI = (TN-TF)/(TN+TF), PI = TN/(TN+TF).

### Cue dependent fear conditioning (CDFC)

Cue dependent fear conditioning test determines long term associative memory involving hippocampal-amygdala axis, where animals learn to associate between an unconditional aversive stimulus of foot shock (0.4 mA, 1 sec for mouse) with a conditional neutral stimulus of light and sound (80 dB, 2500 Hz). Rodents express their fear towards an aversive stimulus by freezing with complete loss of movement except for breathing.

On day 1 or acquisition day, mice were placed into the conditioning chamber 30 mins after any sort of treatment. The test was performed with 2 mins habituation period followed by four consecutive presentations of light and sound for 29 sec each terminating with foot shock. Next it was followed by a 60-s intertrial interval followed by a 2 mins period without stimuli (8 min total). On the probe day the context of the chamber was altered with the addition of white paper covering the walls and vanilla essence. On the probe day, mice were exposed to the same testing paradigm as on conditioning day but in absence of shock stimuli. On both days the freezing % was measured by PACKWIN 2.0 software (Panlab, Barcelona, Spain).

### Statistical Analysis

To test the significant differences among multiple (more than two) groups all data were analyzed using one way ANOVA analysis followed by Tukey’s multiple comparison of means post hoc test by using Graph pad software. For comparison of significant difference between two groups, student’s t-test analysis was used using the same software. Data are presented as mean ± standard error of mean, and p < 0.05 was considered to be statistically significant.

## Results

### Progressive microglial activation is observed in response to Aβ exposure *in vitro* and in 5xFAD mouse models of AD

At first, we wanted to characterise the initiation and progression of microglial activation in 5xFAD mice using 3 different age groups: 2 months, 6 months and 13 months. Homozygous 5xFAD mice which overexpresses human APP gene with 3 mutations (swidish-K670N, M671L; Florida-1716V and London-V717I) and human PS1 gene with 2 mutations (M146L and L286V), accumulate Aβ as early as 2 months of age (Oakley et al., 2006). However, the memory and cognitive decline starts around 6 months of age (Oakley et al., 2006; Ohno et al., 2006).

We analysed microglial activation by morphological alterations such as increase in size and increased expression of cell surface marker, Iba1 in response to deposited Aβ plaque in the CA1 region of hippocampus of mice brain by immunohistochemistry (IHC). Confocal microscopy after co-immunolabelling of 20 μm thick PFA fixed brain sections with anti-Iba1 and anti-Aβ antibody, detected densely stained microglia cells with increased cell body area and perimeter around Aβ deposits in 5xFAD mice brain of all ages when compared with age matched C57BL/6 wild type mice (Fig. 1A-E). Moreover, cell body area, perimeter and Iba1 expressions were found to be significantly higher in 6 months and 12 months old compared to 2 months old 5xFAD mice brains and this data set positively correlated with Aβ plaque number which also increased with age (Fig 1. A, B, F-H). Having observed the temporal profile of Aβ induced microglial activation *in vivo*, we were intrigued to understand the time kinetics of microglial activation in response to Aβ in primary culture system. For this study, we treated primary microglia cells isolated from neonatal rat pups with 1.5μM of Aβ_(1-42)_ oligomer and incubated for 0-24h (Fig. 2A). Confocal images after immunostaining with anti-Iba 1 antibody showed an increase in microglial activation as observed by increased cell body area and elevated Iba1 intensity from 1h till 24h of Aβ exposure (Fig 2. B-D).

**Fig. 1.**
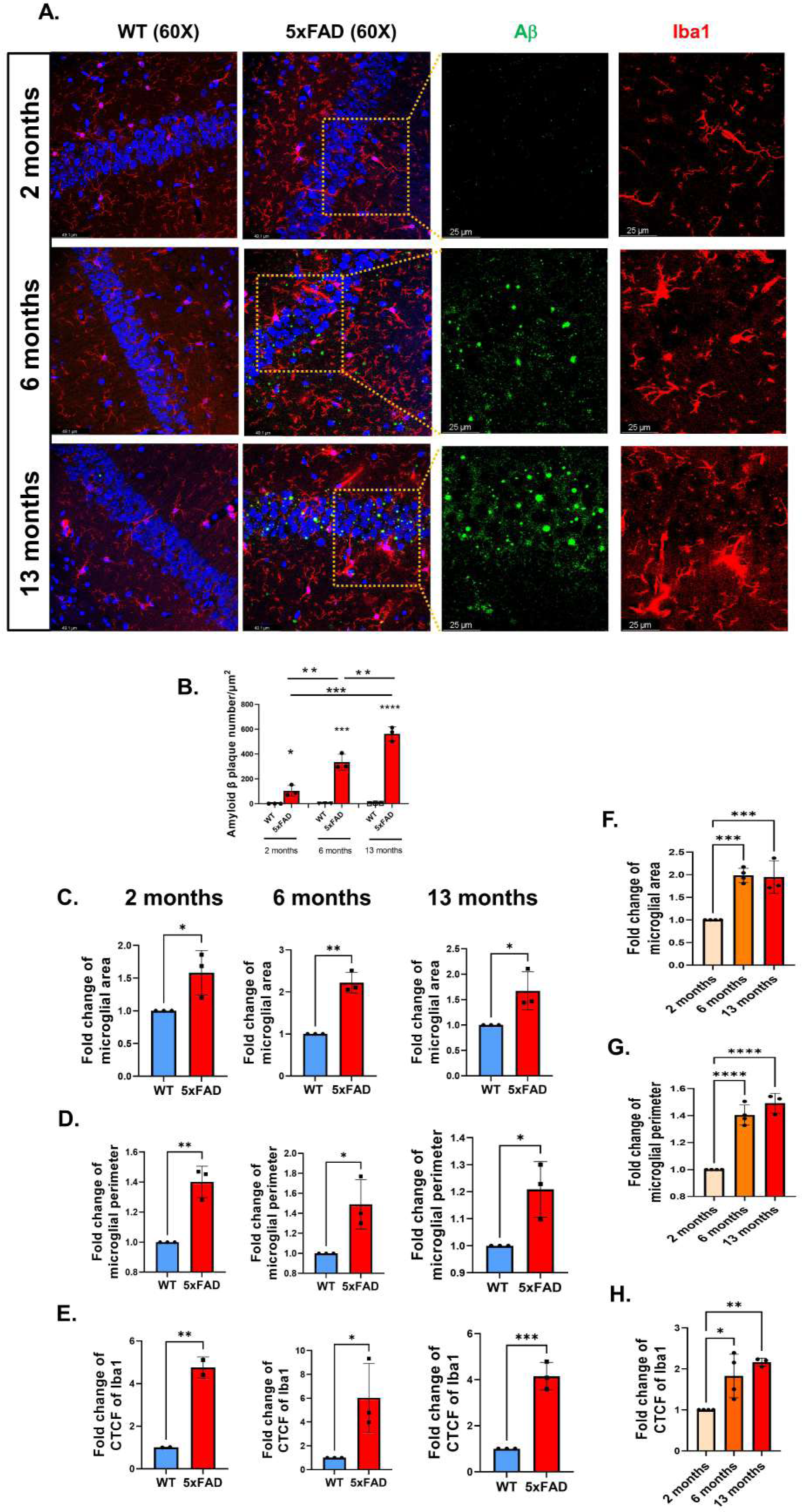
Temporal profile of microglial activation in 5xFAD and C57BL/6 mice brain. **(A)** Representative confocal images of 20μm brain sections of 5xFAD and age matched C57BL/6 mice at 2 months, 6 months and 13 months of age. Hippocampal sections were co-immunolabeled with Iba1 (red) and Aβ (green). Graphical representation of **(B)** Aβ plaque number**, (C, F)** fold change of plaque associated microglial cell body area, **(D, G),** microglial cell body perimeter and **(E, H)** corrected total cell fluorescence (CTCF) of microglial activation marker Iba1; (n=3). Each point of the Scatter plots with bar represents one animal. Data shown mean ± SEM with *P<0.05, **P<0.01, ***P<0.001 and ****P<0.0001.

**Fig. 2.**
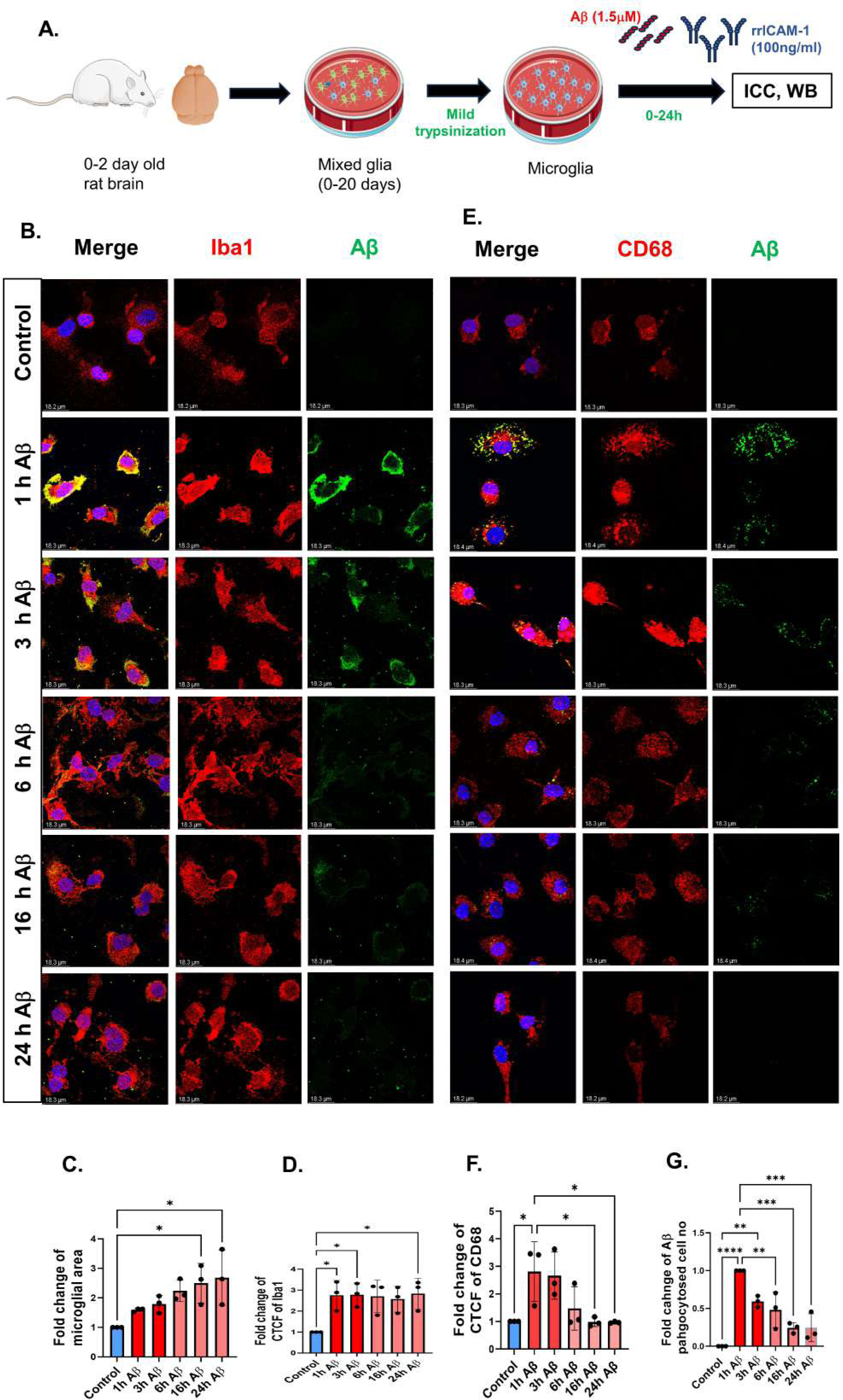
Aβ induces differential activation and phagocytic induction of primary microglia cells. **(A)** Experimental design. **(B)** Representative confocal images of primary microglia cells treated with 1.5μM of Aβ for indicated time periods. Cells were fixed and immunolabeled with anti-Aβ and anti-Iba1 antibodies. Nuclei were stained with Hoechst. Graphical representation of **(C)** fold change of cell body area and **(D)** corrected total cell fluorescence (CTCF) of Iba1. **(E)** Representative confocal images of primary microglia cells treated with 1.5μM of Aβ for time points as indicated and immunostained to visualize phagocytic marker CD68 (red) and Aβ (green). Graphical representations of fold change of CTCF of CD68 **(F)** and Aβ phagocytosed cell number **(G)**. Data were calculated from 3 independent culture experiments. Each point of the Scatter plots with bar represents one independent culture. Data shown mean ± SEM with *P<0.05, **P<0.01, ***P<0.001 and ****P<0.0001.

As microglial activation is often characterised by its ability to phagocytose Aβ, we co-immunolabelled primary microglia cells under same conditions with anti-CD68 antibody, the phagocytic receptor protein and anti-Aβ antibody. Confocal microscope images showed an overall increase in microglial uptake of Aβ as observed by increased number of Aβ engulfing cells along with elevated expression of CD68 within 1h of Aβ treatment which gradually lowered from 6h onwards (Fig 2E-G). These results indicate that Aβ not only induces microglial activation but also its phagocytic ability which may be an essential act of defence.

### Aβ induces microglial inflammation by activating ERK-STAT3 pathway

As microglia are the major effector cells responsible for Aβ induced neuroinflammation, we next wanted to characterise oligomeric Aβ_(1-42)_ induced microglial activation based on their production of pro-inflammatory proteins. Primary microglia cells isolated from neonatal rat pups were subjected to Aβ treatment for 0-24h. Western blot results showed significant increase in protein levels of ROS producing enzyme iNOS, inflammation inducing transcription factor NFκB and pro-inflammatory cytokines such as TNF-α and IL-1β as early as 1-3h of treatment which significantly lowered from 6h onwards (Fig. 3A-F).

**Fig. 3.**
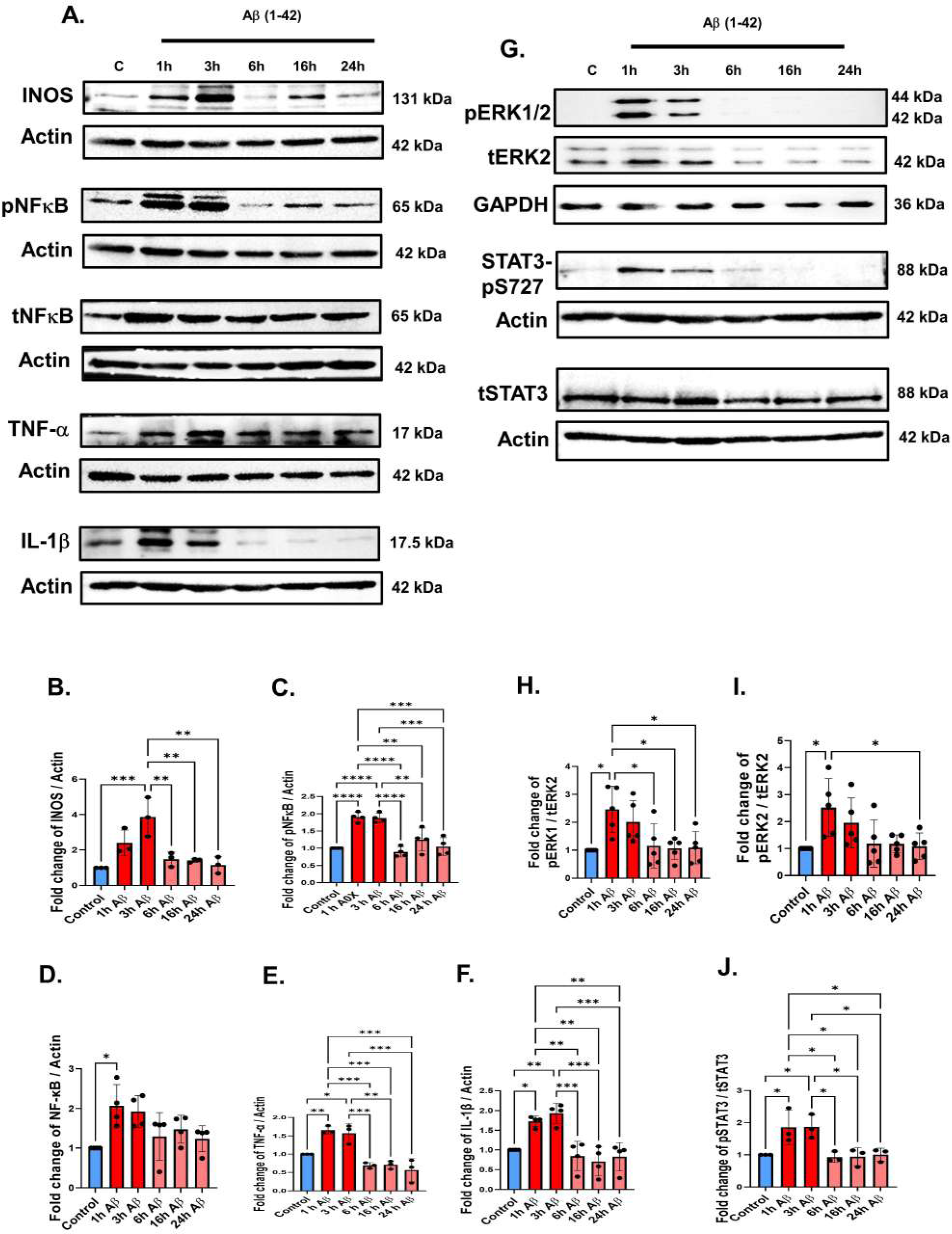
Aβ induces differential expression of inflammatory proteins and ERK, STAT3 phosphorylation. **(A)** Representative western blots showing endogenous levels of different inflammatory proteins from microglia cells after 1.5 μM Aβ treatment for indicated time periods. Graphical representations of fold change of **(B)** INOS, **(C)** pNFκB **(D)** tNFκB **(E)** TNF-α, and **(F)** IL-1β. Data were normalized with actin. **(G)** Representative immunoblots for pERK1/2 and pSTAT3-S727 with corresponding total proteins along with GAPDH and actin respectively from primary microglia cells treated with 1.5 μM Aβ. Graphical representations of fold change of **(H)** pERK1, **(I)** pERK2, and **(J)** pSTAT3-S727. Data were normalized with respective total proteins, (n ≥ 3). Each point of the Scatter plots with bar represents one independent culture. Data shown mean ± SEM with *P<0.05, **P<0.01, ***P<0.001 and ****P<0.0001.

Microglia are one of the major secretory cells of CNS and their functional activation is often reflected in their sceretome profile. To determine if microglial inflammatory activation is mirrored in its secretion, we used an antibody-based detection system that identified differential secretion of proteins in the supernatant of primary microglia cells treated with Aβ for 1 h (Additional file 2). Among the elevated proteins identified in 1h Aβ-treated supernatant, majority of them were found to be pro-inflammatory in nature such as CCL2, CCL5, CCL20, GM-CSF, RGMA, SCF, Tim1, Wisp1; some were phagocytosis inducers like RAGE, galectin, CCL2, SCF, VCAM-1, GM-CSF etc (Additional file 2-3) and these data further support our previous observations.

Since we have established the kinetics of Aβ induced microglial activation we were curious to investigate the mechanism lying behind it. As already discussed, ERK-STAT3 pathway has been closely linked with glial activation and inflammation. Taking cue from previous reports (Huiliang et al., 2021), we next investigated the involvement of ERK in microglial inflammation. A two-fold increase in phosphorylation of ERK1/2 and STAT3 at S727 was observed in response to oligomeric Aβ_(1-42)_ within 1-3 h treatment which significantly lowered from 6h and comparable to the control (Fig.3 G-J). This data was further validated by confocal microscopy where significant increase in intensity of p-ERK was observed after 1-3h of Aβ treatment which went down with increasing exposure time (Fig.4 A-B). Moreover, from the secretome profile we identified a number of upregulated proteins like CCL2, CCL3, CCL5, IGFBP3, SCF whose secretion and function are reported to be regulated by ERK-STAT3 pathway (Additional file 2-3).

**Fig. 4.**
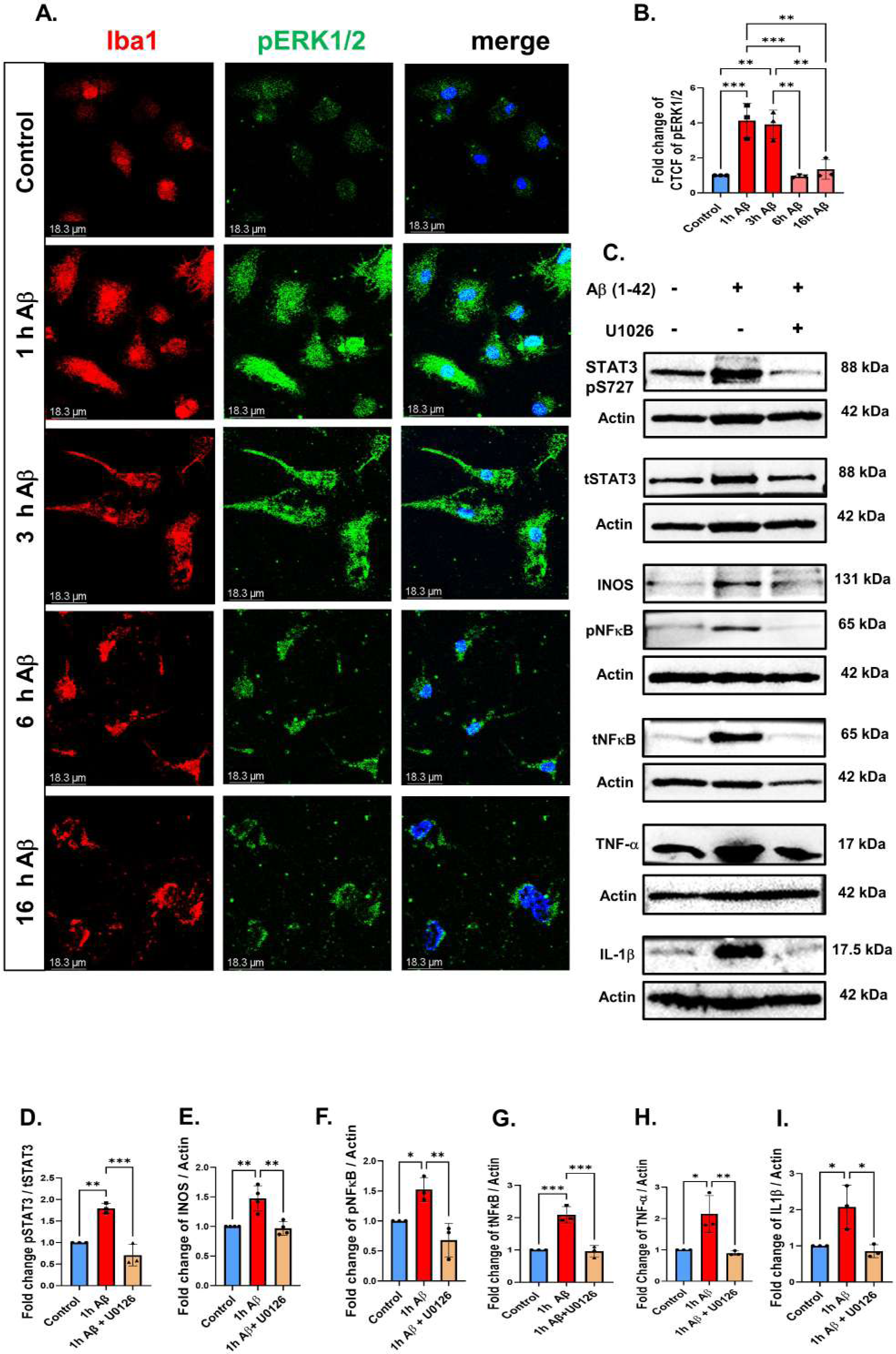
ERK-STAT3 pathway is indispensable for Aβ-induced microglial inflammation. **(A)** Representative confocal images of primary microglia cells treated with 1.5 μM Aβ for indicated time points and stained with Iba1 (red) and pERK1/2 (green) antibodies. **(B)** Graphical representation of fold change of differential intensity of pERK, (**C)** Western blots showing differential levels of pSTAT3, tSTAT3, INOS, pNFκB, tNFκB, TNF-α, IL-1β in cultured primary microglia cells treated with 1.5μM Aβ for 1h in presence or absence of ERK phosphorylation inhibitor U0126. Densitometric analysis and calculation of fold change of **(D)** pSTAT3 / tSTAT3 **(E)** INOS **(F)** pNFκB **(G)** tNFκB **(H)** TNF-α **(I)** IL-1β. Data were calculated from 3 independent culture experiments. Each point of the Scatter plots with bar represents one independent culture. Data shown mean ± SEM with *P<0.05, **P<0.01, and ***P<0.001.

To further validate the essential involvement of ERK-STAT3 pathway in microglial inflammation, we treated primary microglia cells with oligomeric Aβ_(1-42)_ in presence or absence of ERK phosphorylation inhibitor, U0126 (10μM) for 1 h. As expected, in absence of active ERK the level of pSTAT3 at serine 727 was found to be low in presence of U1026. Interestingly the results further showed that in presence of U0126, Aβ mediated induction of inflammatory proteins such as iNOS, pNFκB, tNFκB, TNF-α, IL-1β were significantly reduced (Fig. 4C,D-I). These results indicate the quintessential role of ERK-STAT3 signalling in Aβ-mediated microglial inflammation and this pathway may act as a potential target to reduce microglial activation and associated neuroinflammation.

### ICAM-1 reduces aberrant microgliosis and improves microglial phagocytic ability to engulf Aβ

We have previously established ICAM-1 as neuroprotective molecule but how ICAM-1 was providing neuroprotection was not very clear from our previous observation (Guha et al., 2022). Moreover, Kim et al (Kim et al., 2012) showed the involvement of ICAM-1 in induction of microglial Aβ degrading enzyme neprilysin. Microglia is also reported to be a source of ICAM-1(Hailer et al., 1997). Therefore, we first investigated the temporal profile of both endogenous and secretory ICAM-1 in primary microglia cells after oligomeric Aβ_(1-42)_treatment to find a link between microglial activation and ICAM-1. We observed a significant increase of soluble ICAM-1 level in the supernatant from microglia cells after 6h incubation with oligomeric Aβ_(1-42)_ and this ELISA data negatively correlated with our previously observed temporal profile of microglial inflammation (Fig. 5A). However, the endogenous levels of ICAM-1 decreased from 1h onwards further indicating reduced intracellular reserve due to increased secretion (Fig. 5B,C). Moreover, LFA-1, the cognate receptor of ICAM-1 was found to be expressed in primary microglia cells (Fig. 5D). Taking together we hypothesized that ICAM-1 may have a very important role in Aβ-induced microgliosis.

**Fig. 5.**
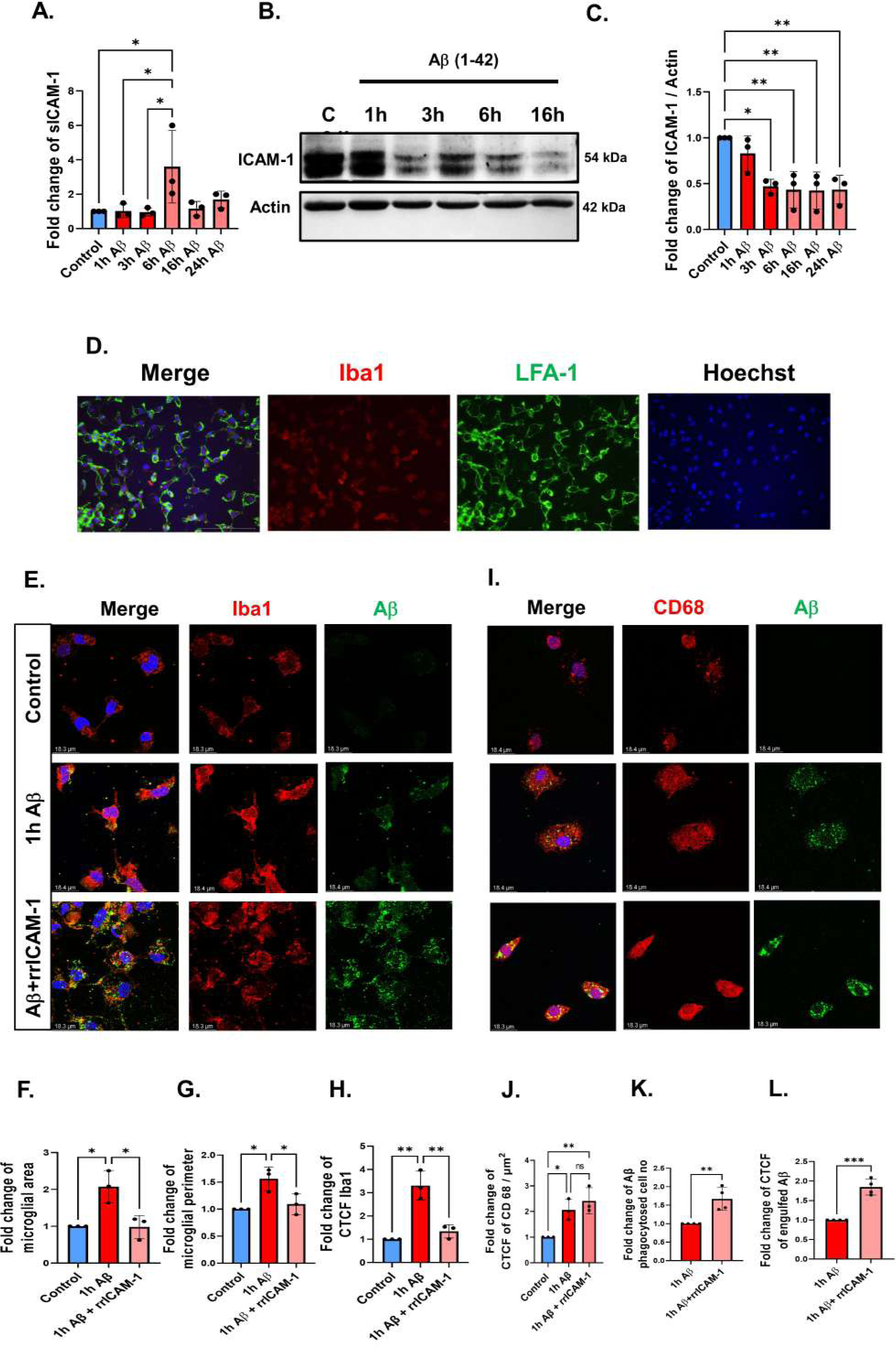
Effect of exogeneous ICAM-1 treatment on microglial activation and phagocytic induction *in vitro*. **(A)** Primary microglia cells were treated with 1.5μM of Aβ in presence or absence of 100ng/ml rrICAM-1 and incubated for indicated time points. The supernatants were collected and subjected to ELISA and fold change of soluble ICAM-1 concentration is graphically represented. **(B)** Western blots depicting endogenous ICAM-1 levels along with actin in response to Aβ treatment. **(C)** Graphical representation of endogenous ICAM-1 levels under indicated conditions. **(D)** Fluorescent microscope images depicting the expression of ICAM-1 receptor LFA-1 in cultured primary microglia cells. **(E)** Primary microglia cells were treated with 1.5μM of Aβ in presence or absence of 100ng/ml rrICAM-1 and incubated for 1h. Cells were fixed and co immunolabeled for Iba1 (Red), Aβ (green) and Hoechst (blue) for nuclei. Graphical representation of fold change of microglial **(F)** cell body area, **(G)** perimeter **(H)** corrected total cell fluorescence (CTCF) of Iba1. **(I)** Primary microglia cells under similar conditions were co-immunolabeled for CD68 (red), Aβ (green) and Hoechst (blue) for nuclei. Graphical representation of **(J)** fold change of CTCF of CD68, **(K)** Aβ-phagocytosed cell number, (**L)** fold change of CTCF of internalized Aβ. As control cells were not treated with Aβ, we excluded control groups from Aβ calculation. Data were calculated from 3 independent culture experiments. Each point of the Scatter plots with bar represents one independent culture. Data shown mean ± SEM with *P<0.05, **P<0.01, and ***P<0.001; ns, not significant.

To determine the effect of ICAM-1 on microglial activation, we treated primary microglia cells with 1.5μM of oligomeric Aβ_(1-42)_ for 1h in presence or absence of rrICAM-1 at a dose of 100ng/ml. Performing an immunocytochemistry in rrICAM-1 treated cells we observed a significant reduction in microglial cell body area, cell body perimeter and expression of activation marker Iba1 compared to cells under only Aβ treatment (Fig. 5E-H).

We have also observed increase but not significant in the expression of phagocytic receptor CD68 in cells treated with ICAM-1 with Aβ compared to cells treated with only Aβ. But ICAM-1 improved Aβ engulfment as observed by increased % of cells engulfing Aβ along with increased internalisation of Aβ indicating a role of ICAM-1 in ameliorating Aβ phagocytosis (Fig. 5I-L).

To further validate the effect of rrICAM-1 on Aβ associated microglial activation and phagocytic induction, we intraperitoneally delivered of rrICAM-1 at a dose of 1μg/kg bodyweight in 5xFAD mice every alternate day for 21 days (Fig. 6A). After co-immunolabeling with anti-Iba1 and anti-Aβ antibodies, confocal images showed that ICAM-1 treatment not only reduced Aβ plaque number but also reduced microglial activation as observed by lowered number of plaque associated microglia with reduced cell body area, perimeter and Iba1 expression in CA1 of hippocampus compared to age matched C57BL/6 mice (Fig. 6B-G). Hence, these results further support the importance of ICAM-1 in modifying microglial activation and inducing phagocytic removal of Aβ.

**Fig. 6.**
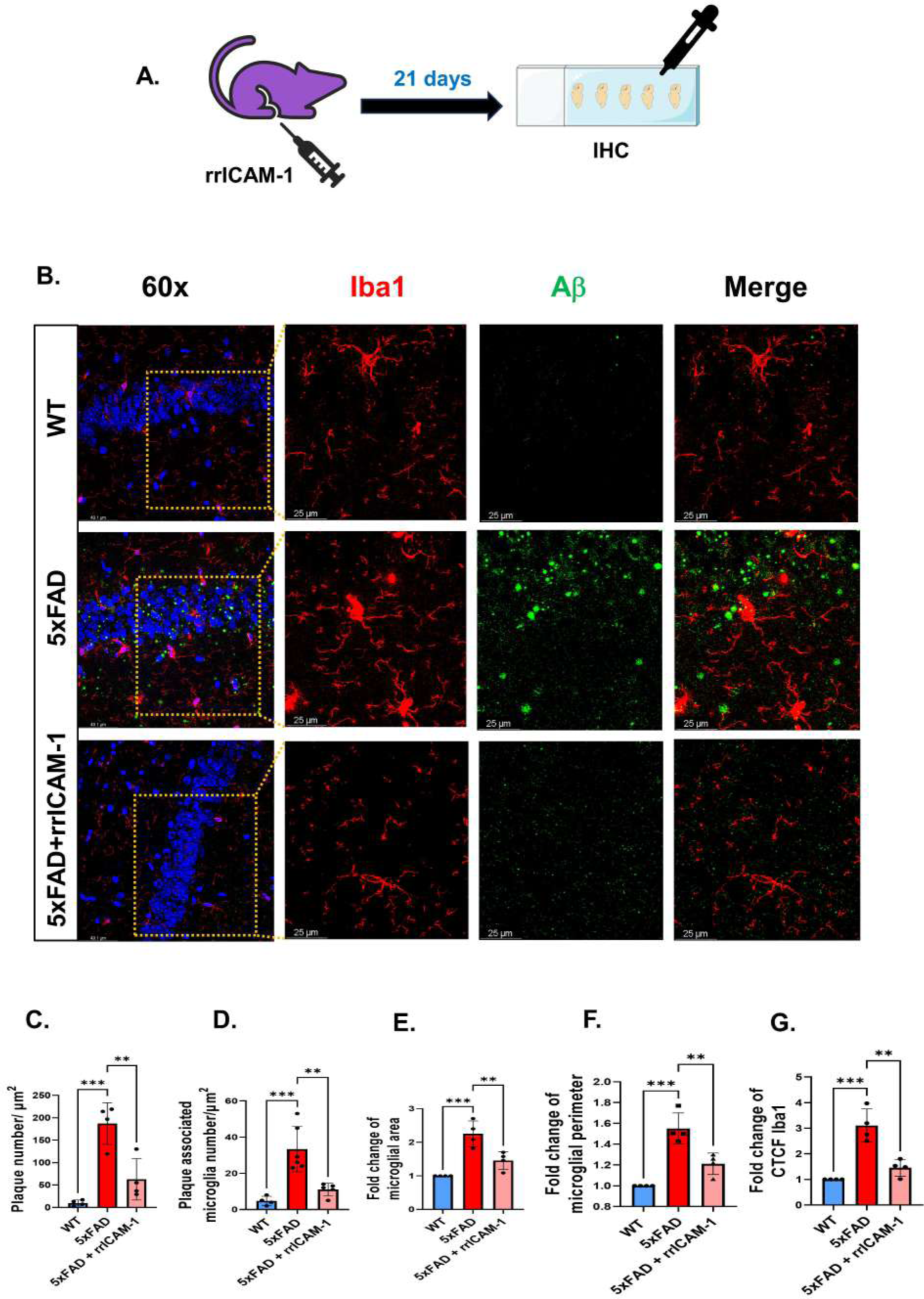
Effect of exogeneous ICAM-1 treatment on microgliosis *in vivo*. **(A)** rrICAM-1 at a dose of 1μg/kg bodyweight was administered intraperitoneally in mice. After fixation, 20μm brain sections were immunolabelled with anti-Iba1 and anti-Aβ antibodies. **(B)** Representative confocal images showing the effect of intraperitoneal ICAM-1 treatment on Aβ plaque (green) and associated microgliosis (red) in the hippocampal CA1 region of 5xFAD compared to age matched C57BL/6 mice. Graphical representation of **(C)** Aβ plaque number, **(D)** plaque associated microglia number, **(E)** fold change of microglial cell body area, **(F)** cell body perimeter, and **(G)** corrected total cell fluorescence (CTCF) of Iba1. Each point of the Scatter plots with bar represents one animal. Data shown mean ± SEM with *P<0.05, **P<0.01, and ***P<0.001.

### ICAM-1 reduces microglial inflammatory activation by inhibiting ERK-STAT3 signalling pathway

To investigate the effect of ICAM-1 on Aβ -induced production of different inflammatory proteins. We treated primary microglia cells with 1.5μM oligomeric Aβ_(1-42)_ in presence or absence of rrICAM-1 at a dose of 100 ng/ml and incubated for 0-3h. The western blot results showed an almost two-fold decrease in the expression of pro-inflammatory cytokines TNF-α and IL-1β levels in presence of ICAM-1 compared to cells treated with Aβ only. (Fig 7A, B-C). However, no significant reduction of iNOS was observed by ICAM-1 treatment. NFκB, the key transcription factor, regulates the production of different pro-inflammatory proteins such as TNF-α, IL-1β etc. by innate immune cells like microglia. Previously we have identified a prominent role of ICAM-1 in inhibition of NFκB levels in 5xFAD mice brain (Guha et al., 2022). This information led us to investigate the effect of ICAM-1 on the functional activation of NFκB in primary microglia cells. The results showed a significant inhibition on the level of phosphorylated and activated p65 subunit of NFκB along with reduction in the expression of the total protein (Fig. 7A, D-E). These findings suggest a prominent anti-inflammatory effect of ICAM-1 on Aβ-mediated microglial inflammation.

**Fig. 7.**
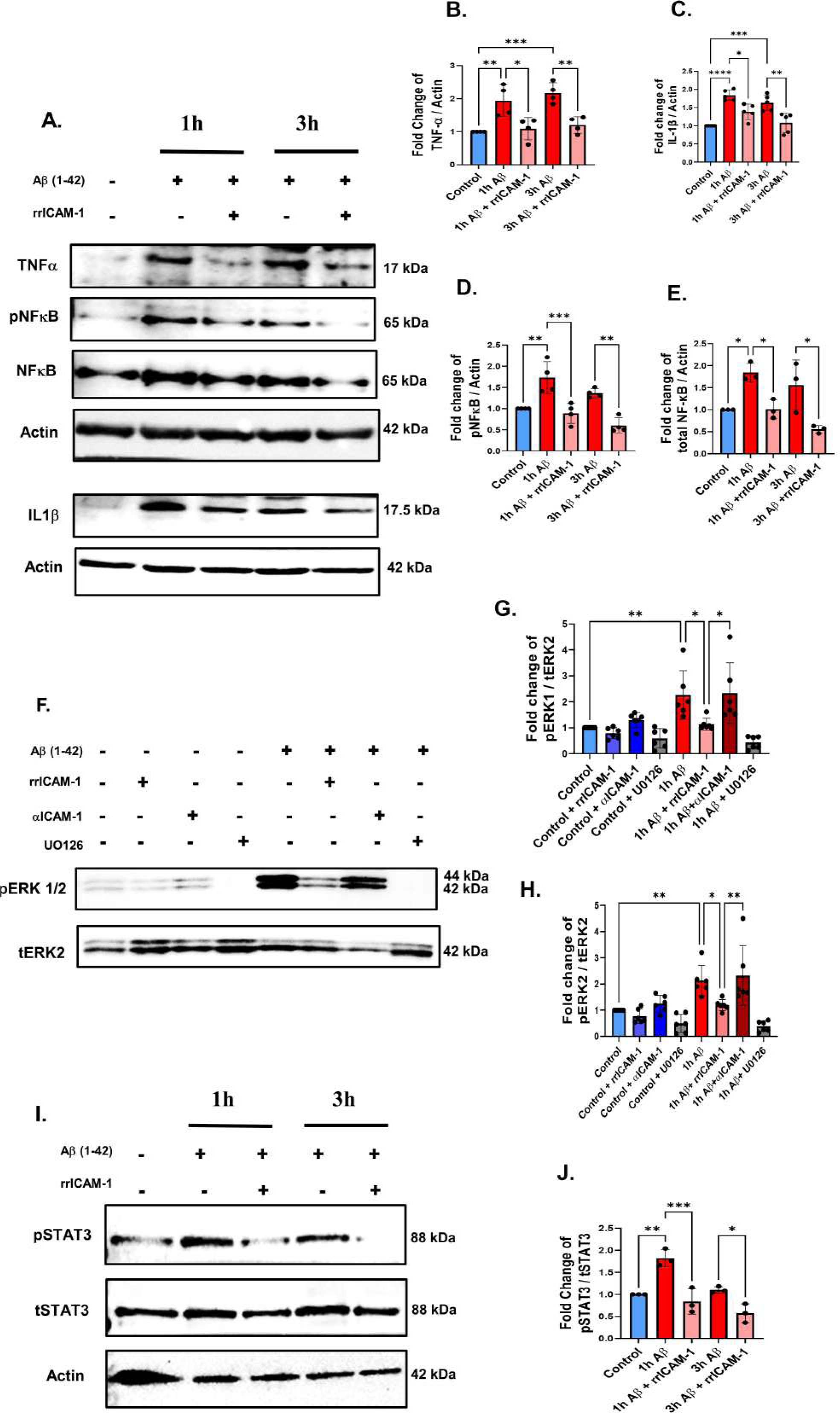
Effect of exogeneous ICAM-1 treatment on ERK mediated microglial inflammation. **(A)** Primary microglia cells were treated with 1.5μM of Aβ in presence or absence of 100ng/ml rrICAM-1 and incubated for indicated time points. Cell lysates were subjected to western blot to assess the levels of inflammatory proteins like TNF-α, pNFκB, tNFκB, IL-1β. Graphical representation of fold change of **(B)** TNF-α, **(C)** IL-1β, **(D)** pNFκB, and **(E)** tNFκB. Data were normalized with actin, (n ≥ 3). **(F)** Primary microglia cells were treated with 1.5μM of Aβ,100ng/ml rrICAM-1, 4μg/ml ICAM-1 neutralizing antibody and 10 μM of U0126 as indicated and incubated for 1h. Cell lysates were used for western blot to check the levels of pERK1/2. Graphical representation of fold change of **(G)** pERK1 and **(H)** pERK2. Data were normalized with total protein, (n=3). **(I)** The representative immunoblots for pSTAT3 and total STAT3 along with actin in presence or absence of rrICAM-1 after Aβ treatment. **(J)** Graphical representation of fold change of pSTAT3 normalized with tSTAT3, (n=3). Each point of the Scatter plots with bar represents one independent culture. Data shown mean ± SEM with *P<0.05, **P<0.01, ***P<0.001 and ****P<0.0001.

Since we have already shown that ERK-STAT3 signalling pathway plays a significant role in microglial inflammation, we next investigated the effect of ICAM-1 on ERK-STAT3 signalling. Primary microglia cells were treated with oligomeric Aβ_(1-42)_ along with rrICAM-1 and ERK phosphorylation inhibitor U0126 at a dose of 10μM. Results showed significant reduction of phosphorylated ERK1/2 levels in presence of ICAM-1 indicating inhibited activation when compared with Aβ only. As microglia is also a source of secretory ICAM-1 we neutralised the endogenously secreted ICAM-1 present in the media by anti-ICAM-1 antibody to prevent the autocrine action of ICAM-1 on microglia. We have observed restoration of phosphorylated ERK1/2 levels in the absence of exogenous or endogenously secreted ICAM-1 (Fig. 7F-H). Moreover ICAM-1 mediated inhibition of ERK1/2 phosphorylation resulted in reduced phosphorylation of STAT3 at serine 727 (Fig. 7I-J). These results clearly point towards the role of ICAM-1 in targeting ERK-STAT3 pathway to modulate Aβ mediated microglial inflammation

### ICAM-1 mediated cognitive improvement and restoration of synaptic protein expression require microglial LFA-1

Since we have already established the inhibitory effect of ICAM-1 on Aβ associated microgliosis, we next investigated the importance of ICAM-1-LFA-1 interaction in ICAM-1 function. Studies indicate that receptor LFA-1 is found to be expressed majorly in microglia cells apart from short lived leukocytes that migrate through leaky blood brain barrier in CNS of AD brain (Hailer et al., 1997). LFA-1 is a heterodimeric integrin (α1β2) with 190 amino acid long ICAM-1 binding domain called inserted domain or I domain present on the α1 integrin (Shimoka et al 2002).

Lifitegrast, a tetrahydroisoquinoline derivative, competitively binds to the I domain of αl subunit and blocks or inhibits ICAM-1 binding. To block the interaction between ICAM-1 and LFA-1 in microglia of brain without affecting the migration of peripheral leukocytes through blood brain barrier route, we intranasally delivered Lifitegrast at a dose of 10 μM 2h prior to rrICAM-1 intraperitoneal injection on every alternate day for 21 days before sacrifice (Fig. 8A). To investigate the effect of lifitegrast on ICAM-1 mediated restoration of microglial homeostasis, we co-immunolabeled brain sections of 5xFAD mice treated with or without lifitegrast along with ICAM-1 administration with anti-Iba1 and anti-Aβ antibody. Confocal imaging analysis data showed that ICAM-1 mediated reduction of Aβ plaque and associated microgliosis indicated by plaque associated microglia number, cell body area, perimeter and Iba1 expression were partially reduced by lifitegrast pre-treatment (Fig. 8B-F).

**Fig. 8.**
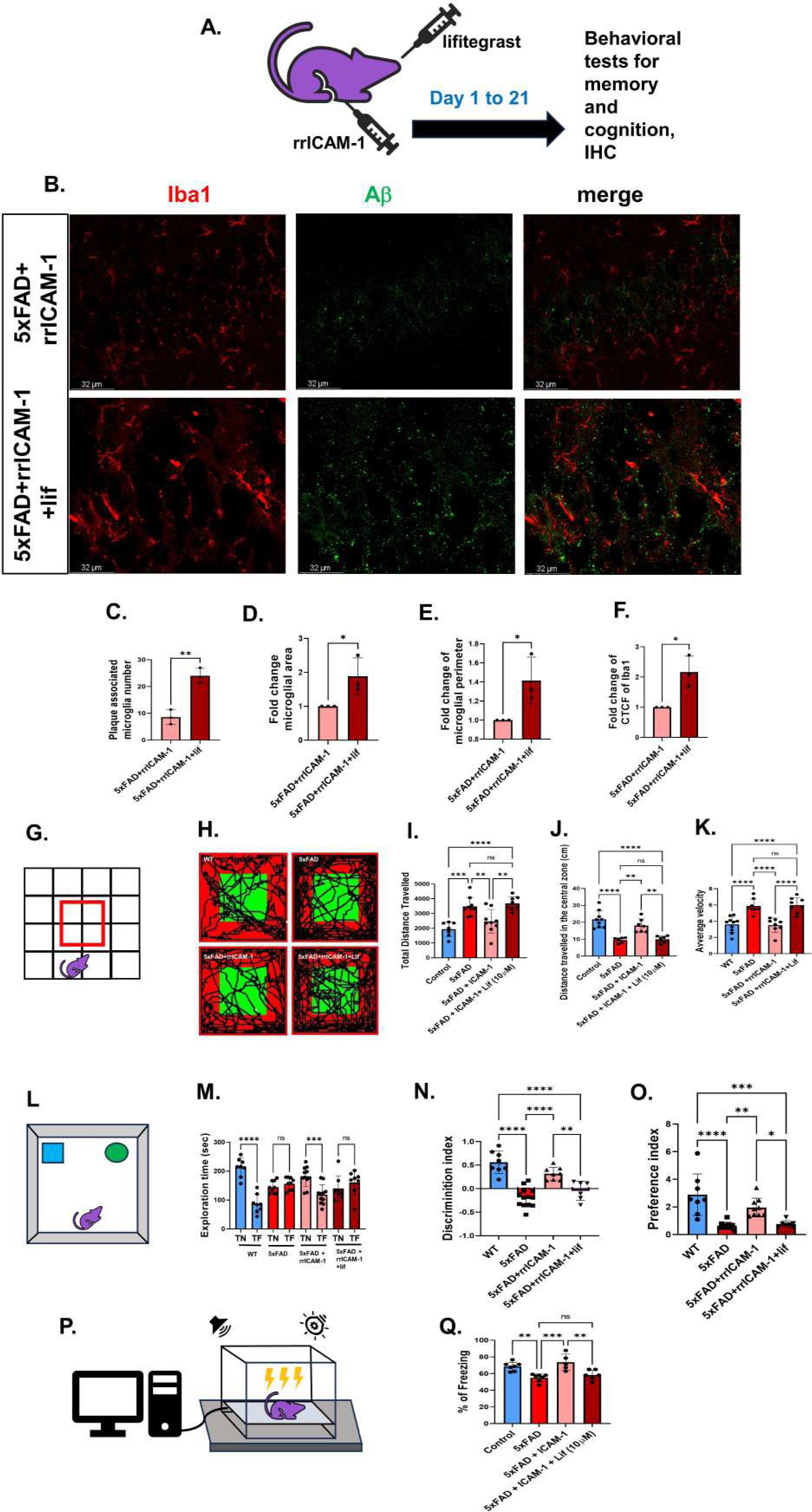
Effect of lifitegrast on ICAM-1 mediated cognitive improvement. **(A)** Lifitegrast at a dose of 10μM has been administered intranasally 2h before rrICAM-1 treatment and a battery of behavioral tests was performed with four groups of mice - WT, 5xFAD, 5xFAD+rrICAM-1 and 5xFAD+rrICAM-1+ Lifitegrast (lif). **(B)** Representative confocal images showing Aβ plaque (green) and associated Iba1 positive microglia cells (red) in CA1 of hippocampus of 5xFAD mice brain with or without lifitegrast administration before rrICAM-1. Graphical representation of **(C)** plaque associated microglia number, **(D)** fold change of microglial cell body area, **(E)** cell body perimeter, and **(F)** corrected total cell fluorescence of Iba1, (n=3). **(G)** Experimental set up for open field test. **(H)** Representative infrared image of path travelled by mice from all 4 groups. Graphical representations of **(I)** average distance travelled by the animals in the open field arena in 10 minutes of time, **(J)** average distance travelled by the animals in the central zone, and **(K)** mean velocity of animals. **(L)** Experimental image of novel object recognition on day 2 with one novel and one familiar object. Graphical representations of **(M)** exploration time near novel (N) or familiar (F) object by animals in sec, **(N)** discrimination index, and **(O)** preference index, (n≥ 6). **(P)** Experimental representation of cue dependent fear conditioning test. **(Q.)** Graphical representation of percentage freezing of mice on day 2. Each point of the Scatter plots with bar represents one animal. Data shown mean ± SEM with *P<0.05, **P<0.01, ***P<0.001 and ****P<0.0001; ns, not significant.

Next, we investigated the importance of this interaction for ICAM-1 mediated cognitive improvement. From day 15^th^ of treatment, we started to perform a battery of behavioural tests such as open field test (OFT) for general locomotion and anxiety, novel object recognition (NOR) for recognition memory and cue dependent fear conditioning (CDFC) for associative memory with mice divided in the following 4 groups C57BL/6 (wild type), 5xFAD, 5xFAD+rrICAM-1, 5xFAD+rrICAM-1+lifitegrast.

In OFT, within 10 mins of time, both 5xFAD mice and mice, where LFA-1-ICAM-1 binding was interfered by lifitegrast treatment, showed hyper locomotion with higher average distance (in cm) travelled, with higher mean velocity and increased anxiety as determined by avoidance of central zone exploration compared to 5xFAD mice which received rrICAM-1 injection without lifitegrast and wild type animals (Fig. 8G-K).

Lifitegrast was also found to block ICAM-1 mediated restoration of novel object recognition memory as observed by reduced exploration time with the novel object on day 2 which was quite comparable to 5xFAD mice. This was further supported by negative value of discrimination index and lower value of preference index which denote lack of ability to discriminate between novel and familiar object when compared with mice where ICAM-1-LFA-1 interaction was not blocked (Fig. 8L-O). Thus ICAM-1-LFA-1 interaction plays important role in amelioration of recognition memory in 5xFAD mice.

In CDFC test, after a training with cued adverse stimulation on day one, % of freezing within 8 mins period was recorded with 4 simultaneous sessions of 2 different cues without adverse stimuli on day two. On day two of CDFC, ICAM-1 treated mice were able to associate between the cue and the adverse stimuli as observed by increased % freezing in response to only cue, which were similar to the wild type mice. ICAM-1 mediated associative memory retention could not be observed in 5xFAD mice which did not receive any treatment or mice where ICAM-1-LFA-1 interaction was blocked by lifitegrast treatment. (Fig. 8P-Q).

The functional mechanism behind cognitive improvement lies in synaptic health and it can be measured by the expression of different pre- and post-synaptic proteins. To determine the importance of microglial ICAM-1-LFA-1 interaction on restoration of synaptic health, we performed immunohistochemistry with brain sections of animals from the above mentioned four groups using antibodies against pre-synaptic protein SNAP25 and post-synaptic protein PSD95. Confocal microscope image analysis showed that, protein puncta number in the CA1 of hippocampus, indicative of expression of both the proteins, were found to be restored by ICAM-1 treatment compared to 5xFAD mice (Fig. 9A-C). Microglia plays crucial role in elimination of dying neurons and their synapses during different diseased conditions (Wilton et al., 2019). To further investigate the involvement of microglia in ICAM-1 mediated synapse modification, we analysed the synaptic puncta engulfment by microglia. The data showed that the number of microglia cells with engulfed PSD95 puncta were reduced in 5xFAD mice that received ICAM-1 compared to mice without any treatment (Fig. 9 D-E). Restoration of PSD95 protein expression by ICAM-1 was significantly blocked by lifitegrast pre-treatment. In case of SNAP25, lifitegrast treatment reduced the expression but not significantly (Fig. 9 A-C). Similar observation was made by western blotting where protein expression of SNAP25 and PSD95 were found to be restored by ICAM-1 treatment. In contrast, the restoration of PSD95 protein level by ICAM-1 treatment was significantly blocked by lifitegrast pre-treatment whereas lifitegrast could not significantly block SNAP25 restoration (Fig.9 F-H). Collectively, these results suggest the importance of microglial LFA-1 in ICAM-1 mediated reduction of Aβ associated microgliosis and inflammation, and that reduced amyloid pathology improves cognition and memory by restoring synapses.

**Fig. 9.**
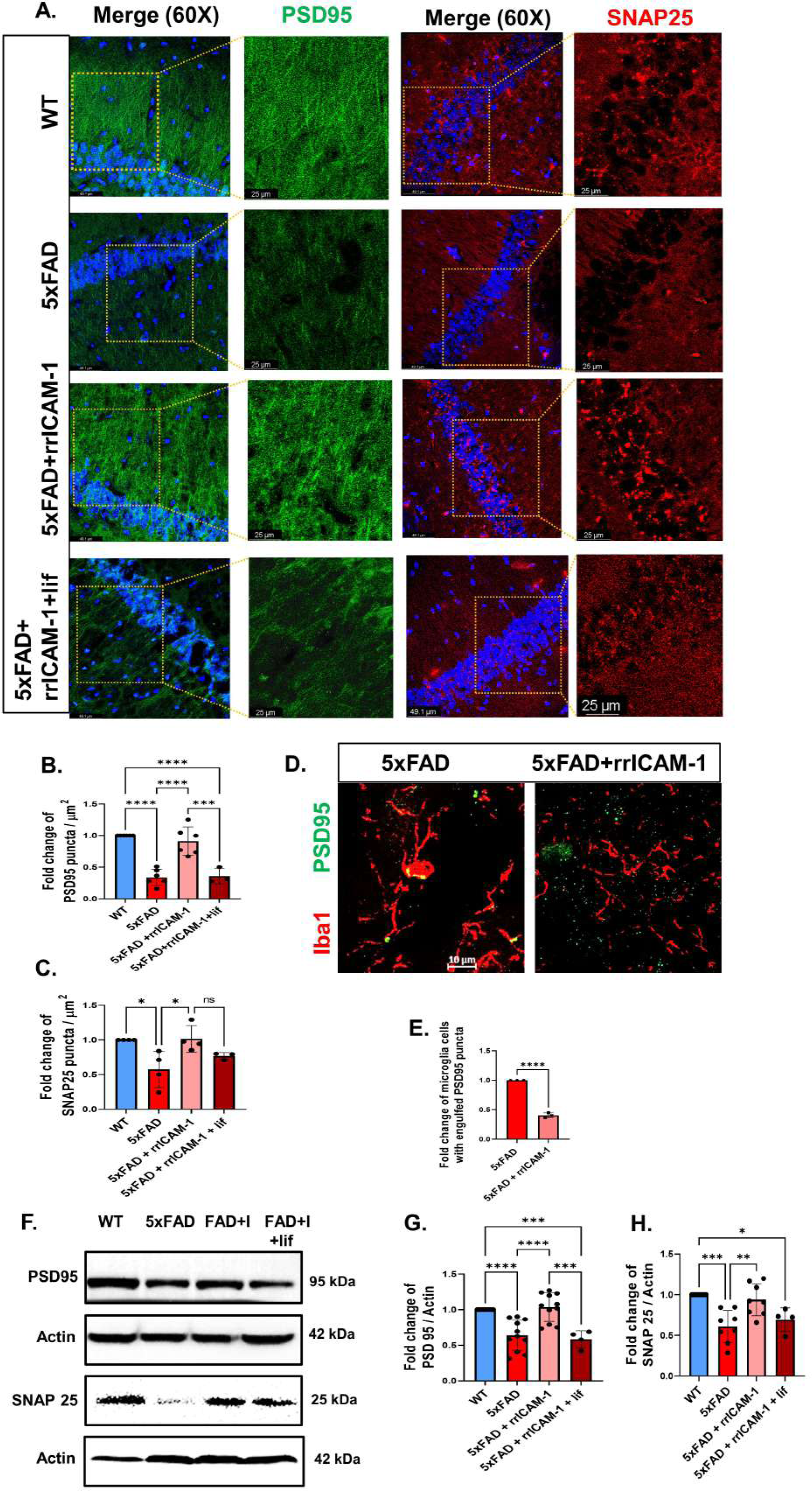
Effect of lifitegrast on ICAM-1 mediated restoration of synaptic protein expressions in 5xFAD mice. Lifitegrast at a dose of 10μM has been administered intranasally before rrICAM-1 treatment at a dose of 1μg/kg bodyweight. After fixation, 20μm brain sections were immunolabelled with post-synaptic PSD95 (green) and pre-synaptic SNAP25 (red) proteins from mice divided in 4 groups WT, 5xFAD, 5xFAD+rrICAM-1 and 5xFAD+rrICAM-1+ Lifitegrast (lif). **(A)** Representative confocal images showing the effect of rrICAM-1 administration with or without lifitegrast pre-treatment, on expression of PSD95 and SNAP25 in the CA1 hippocampal region of 5xFAD mice. **(B-C)** Graphical representations of fold change of SNAP25 and PSD95 puncta number/μm2. **(D)** Representative confocal images showing engulfment of PSD95 puncta (green) by Iba1 positive microglia cells (red) in presence or absence of rrICAM-1 treatment in hippocampal CA1 region of 5xFAD mice brain. **(E**) Graphical representation of fold change of cells with engulfed PSD95 puncta inside. **(F)** After treatment of ICAM-1 in presence or absence of lifitegrast, tissue lysates from WT, 5xFAD, 5xFAD+rrICAM-1 (FAD+I) and 5xFAD+rrICAM-1+ Lifitegrast (FAD+I+lif) were subjected to western blotting to check the endogenous levels of PSD95 and SNAP25. **(G-H)** Graphical representations of fold change PSD95 and SNAP25, (n≥4). Each point of the Scatter plots with bar represents one animal. Data shown mean ± SEM with *P<0.05, **P<0.01, ***P<0.001 and ****P<0.0001; ns, not significant.

## Conclusions

AD being the most common form of dementia currently affects 50 million people worldwide (Taylor et al., 2017). One of the major causes of the disease pathogenesis is Aβ mediated neuroinflammation which lead to AD associated neurodegeneration and cognitive impairment. (Heppner et al., 2015; Hodson, 2018; B. Zhang et al., 2013). Aβ in its soluble oligomeric or fibrillar form is easily degraded by a number of microglia secreted Aβ-degrading enzymes such as Neprilysin, Insulin degrading enzyme, MMP9 and plasminogen. However, its increased production and slow degradation makes microglia overwhelmed and saturated thereby leading to accumulation of excess Aβ in the form of insoluble plaques. (Kato et al., 2022; Leissring et al., 2003). Recently, we have shown that ICAM-1 improves memory and cognitive impairments in 5xFAD mice model of AD (Guha et al 2022). Since LFA-1, the cognate receptor of ICAM-1 is majorly present on microglia cells in CNS, we investigated the role of ICAM-1 on microglia mediated Aβ clearance and cognitive function.

Our experimental findings reveal the complex kinetics of microglial activation and inflammation in response to Aβ in both mice brain hippocampus and primary microglia culture. We found a 2-fold increase of microglial activation marker Iba1 between 2 months and 6 months age linking microglial activation with cognitive impairment that starts around 6 months of age of 5xFAD mice as reported earlier (Oakley et al., 2006; Ohno et al., 2006). We have also observed robust microglial activation in culture with increased expression of pro-inflammatory proteins within 1h of Aβ treatment and this induction of inflammation has reduced from 6h onwards indicating the existence of a complex time dependent induction of microglial inflammation in response to Aβ. It is well documented that ERK activation induces downstream inflammatory proteins like TNF-α, NFκB, IL-1β in various diseased conditions (Guha et al., 2001; Huang et al., 2021; Li et al., 2006; Seo et al., 2013; Shi et al., 2002). In this study we have observed a 2-fold increase in both ERK1 and ERK2 phosphorylation along with the expression of inflammatory proteins indicating a possible role of ERK in microglial inflammatory activation. Chen et al in 2021 (Chen et al., 2021) have reported an upregulation of ERK1/2 phosphorylation in 5xFAD mice hippocampal microglia. This led us to investigate the effect of ERK1/2 inhibition on microglial inflammation. We have found that ERK1/2 activation plays an indispensable role in microglial inflammation because microglial inflammatory protein production has been reduced when ERK phosphorylation was inhibited by U0126, a MEK1/2 inhibitor. ERK, the MAPK specifically targets STAT3 and phosphorylates at serine 727which is essential for gliosis (Kang et al., 2012; Ryu et al., 2019; Yu et al., 2015; B. Zhang et al., 2013; Y.-Y. Zhang et al., 2013). Balic et al (Balic et al., 2020) have shown that STAT3 phosphorylation at serine 727 residue is required for its mitochondrial translocation in order to upregulate inflammatory cytokine such as IL-1β production. We have observed here a simultaneous phosphorylation of STAT3 at serine 727 in response to Aβ which was reduced by inhibition of ERK1/2 phosphorylation. These data strongly suggest that ERK1/2-STAT3 pathway is critical for Aβ induced microglial inflammation. Recently a small chemical derivative of a naturally occurring human adrenal sterol β-androstenetriol have been developed that blocks ERK1/2 mediated activation of inflammation in different neurological conditions including AD (Reading et al., 2021). ERK1/2 is ubiquitously present in different cell types in brain and plays important role in learning and memory functions of brain (Hutton et al., 2017). Thus, microglia specific blocking of ERK1/2 could have good therapeutic potential.

In consistence with previous report (Kim et al., 2012), we have already shown that ICAM-1 has a strong Aβ plaque ameliorating ability in 5xFAD mice brain. (Guha, Paidi et al. 2022). As microglial dysfunction is one of the major causes of Aβ plaque accumulation in CNS we hypothesised the involvement of ICAM-1 in augmenting microglial phagocytosis of Aβ. Moreover, ICAM-1, beyond its function in leukocyte migration, has been linked to neuroinflammation in a variety of clinical contexts (Lindsberg et al., 1996; Sheikh et al., 2023). Hence, we investigated the possible mechanism of ICAM-1 function in transforming microglia into an anti-inflammatory, highly phagocytic, disease evading isotype. According to several reports, microglia is one of the major sources of ICAM-1 and by ELISA we have observed ICAM-1 secretion to be negatively correlating with ERK1/2 mediated inflammation indicating the presence of a feedback mechanism between ICAM-1 secretion and microglial inflammation. To investigate further, we exogenously incorporated ICAM-1 along with Aβ and found that it was able to significantly reduce Aβ induced microglial activation as well as levels of pro-inflammatory proteins indicating a potent anti-inflammatory effect of ICAM-1 on Aβ-induced microglial inflammation. Moreover ICAM-1 treatment inhibited Aβ-induced ERK and STAT3 phosphorylation which further supports the anti-inflammatory property of ICAM-1 with ERK as the possible target of inhibition. Interestingly, ICAM-1 treatment was found to induce Aβ engulfment and phagocytosis in primary microglia cells while reducing its activation as indicated by reduced amoeboid morphology and Iba1 expression. This *in vitro* data was supported by our IHC data where we have observed reduced Aβ plaque number and associated microgliosis after intraperitoneal delivery of ICAM-1 in 5xFAD mice. To further validate our observation, before ICAM-1 injection we intranasally delivered lifitegrast, an antagonist of ICAM-1 to block its interaction with microglia rich receptor LFA-1. In presence of lifitegrast we observed partial loss of ICAM-1 mediated improvement in plaque pathology as well as associated microglial activation. Although LFA-1 is the cognate receptor but ICAM-1 can also function by binding to another receptor called MAC-1. This may explain the partial retention of ICAM-1 function even after blocking of ICAM-1 binding site in LFA-1. Therefore, we inferred that ICAM-1 modifies microglia into a beneficial subtype that clears Aβ plaque in one hand and reduces Aβ associated inflammation on the other hand in 5xFAD mice brain as well as in primary culture by binding to its cognate receptor LFA-1 which is majorly expressed in microglia.

Synapse loss and neurodegeneration are two intertwined incidents and the first one often precedes the latter. In AD, Aβ induced neuroinflammation strongly affects synaptic health and associated brain functions (Prieto et al., 2019). During development microglia are seen to be involved in synaptic pruning but excessive removal can lead to the development of different neurological and psychiatric disorders (Wilton et al., 2019). In this study, we show that ICAM-1 treatment improves synaptic health by restoring post-synaptic PSD95 and pre-synaptic SNAP25 protein expressions in 5xFAD mice when compared to untreated mice. Moreover, ICAM-1 treatment reduced microglia mediated engulfment of PSD95 further supporting the effect of ICAM-1 in microglia mediated restoration of synaptic protein expression. This improved synaptic health is reflected in the memory and cognitive improvement by ICAM-1 treatment as observed in 3 different behavioral experiments. This was further validated by intranasal lifitegrast pre-treatment that blocked microglial ICAM-1-LFA-1 interaction, and inhibited ICAM-1-mediated memory and cognitive improvement in 5xFAD mice.

In conclusion, this work has revealed ICAM-1 as an immunomodulator that potentiates microglial clearance of Aβ while reducing its detrimental and harmful inflammatory activation. This eventually improves Aβ plaque pathology and restores synaptic health which is reflected in the ameliorated cognitive functions in 5xFAD mice.

## Declarations

### Ethics approval and consent to participate

All animal studies were carried out in accordance with the guidelines formulated by the Committee for the Purpose of Control and Supervision of Experiments on Animals (Animal Welfare Divisions, Ministry of Environments and Forests, Govt. of India) with approval from the Institutional Animal Ethics Committee (IAEC). We have consulted the ARRIVE guidelines for the relevant aspects of animal studies.

### Consent for publication

<u>Not applicable</u>

### Availability of data and materials

All data used and analysed in the current study are available with the corresponding author upon request.

### Authors’ contributions

S.G. and S.C.B. conceptualised and visualised; S.G. designed, investigated, validated and analysed the study; S.C.B. supervised the project and provided the necessary resources; S.G. performed all biochemical, histological, cell culture, microscopy and animal experiments; S.G. wrote the original draft and S.C.B reviewed and edited the paper.

## Acknowledgements

The work was supported by the Council of Scientific and Industrial Research, Govt. of India. We thank Mr. Sounak Bhattacharya from Central Instrument Facility (CIF), CSIR-IICB, Kolkata for his contributions in Leica Sp8 STED confocal microscopy image acquisition.

---

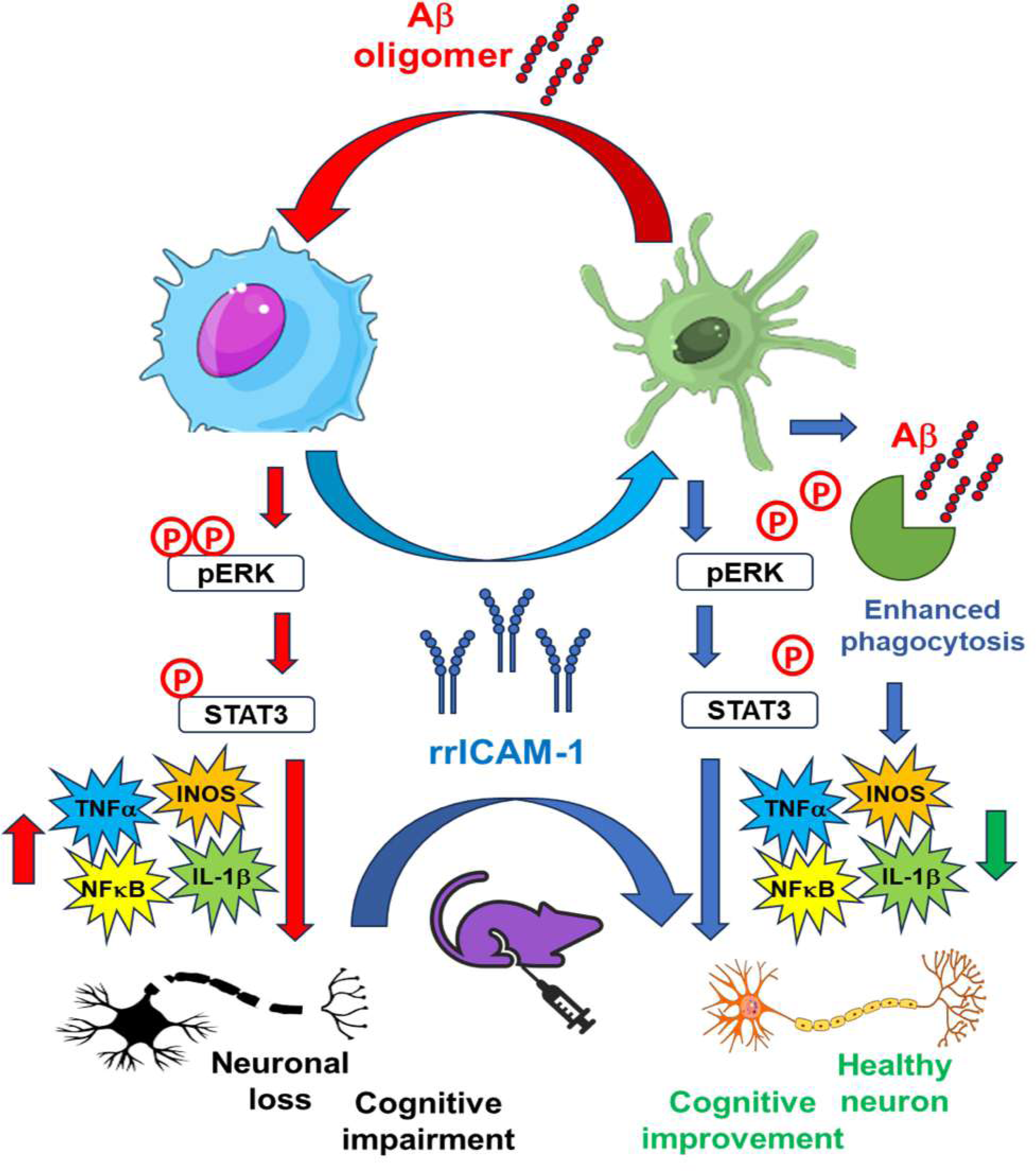

