## Supplementary Materials for "Intercellular adhesion molecule-1 reprograms microglia to improve cognitive functions by inhibiting ERK/STAT3 signalling pathway in a model of Alzheimer’s disease"

**Additional file 1**

| **Antibody** | **Host** | **Application** | **Dilution** | **Source** | **#Cat** |
|---|---|---|---|---|---|
| Actin | rabbit | WB | 1:10000 | Merck | A3854 |
| Amyloid β | rabbit | ICC, IHC | 1:500 | Abcam | Ab201060 |
| Anti-rabbit IgG-HRP linked | Goat | WB | 1:4000 | Cell Signalling Technologies | 7074S |
| Anti-mouse IgG-HRP linked | Horse | WB | 1:4000 | Cell Signalling Technologies | 7076S |
| CD68 | mouse | ICC | 1:50 | Biorad | MCA 1957 |
| GAPDH | rabbit | WB | 1:2000 | Cell Signalling Technologies | 2118S |
| HOECHST | | ICC,IHC | 1:1000 | Invitrogen | H3570 |
| Iba1 | mouse | ICC,IHC | 1: 100 | Merck-millipore | MABN92 |
| ICAM-1 | mouse | WB | 1:500 | Novus | NBP2-22541 |
| IL-1β | rabbit | WB | 1:500 | Abclonal | A1112 |
| INOS | rabbit | WB | 1:500 | Novus | NB300-605 |
| LFA-1 | | ICC | 1:200 | Abcam | AB 13219 |
| pERK (T202/Y204) | rabbit | WB,ICC | 1:1000 | Cell Signalling Technologies | 4377S |
| pNFκB | rabbit | WB | 1:500 | Cell Signalling Technologies | 3033s |
| PSD95 | rabbit | WB, IHC | 1:1000 | Abcam | Ab18258 |
| pSTAT3 | rabbit | WB | 1:1000 | Cell Signalling Technologies | 9134s |
| SNAP25 | rabbit | WB, IHC | 1:1000 | Abcam | Ab108990 |

| | | | | | |
|---|---|---|---|---|---|
| Total ERK | mouse | WB | 1:1000 | Santa Cruz biotechnologies | Sc153 |
| Total NFκB | rabbit | WB | 1:500 | Cell Signalling Technologies | 8242s |
| TNF-α | rabbit | WB | 1:500 | Abclonal | A11534 |
| Total STAT3 | rabbit | WB | 1:500 | Santa Cruz biotechnologies | Sc482 |

**Additional file 1:** List of antibodies. WB=western blot, ICC=immunocytochemistry, IHC=immunohistochemistry

### Additional file 2

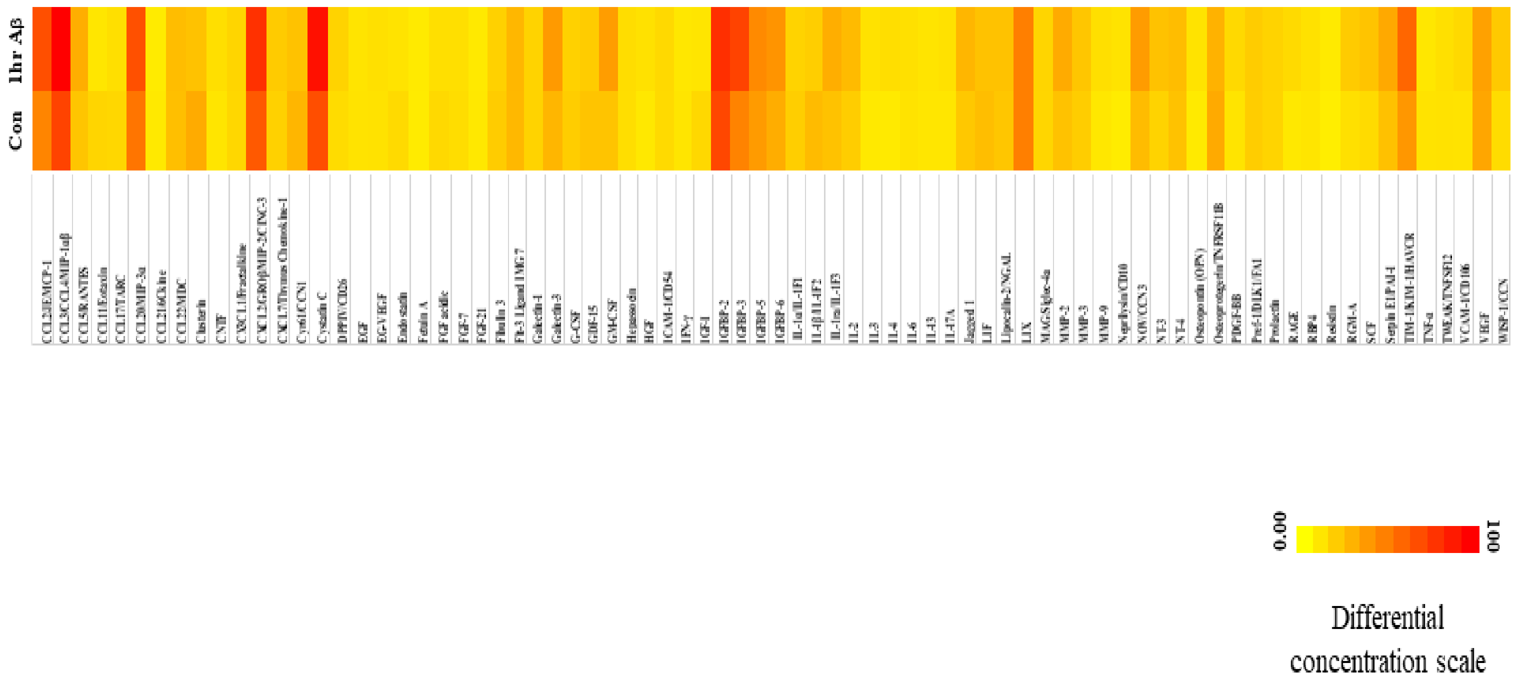


**Additional file 2:** Heat map showing differential secretion of proteins in response to Aβ as observed in cytokine array experiment.

### Additional file 3

| | **Fold change of concentration** | **Function** | **Reference** |
|---|---|---|---|
| ***CCL2/JE/MCP-1*** | 1.439587 | Inflammation, phagocytosis, ERK inducer | (Lee et al., 2021), (Tang & Tsai, 2012) |
| ***CCL3/CCL4/MIP-1α/β*** | 1.350079 | Inflammation, phagocytosis, ERK inducer | (Lindell et al., 2001) (Kodama et al., 2020) |
| **CCL5/RANTES** | 1.39302 | Inflammation, phagocytosis, ERK inducer | (Skuljec et al., 2011) (H. Wang et al., 2023) (Kodama et al., 2020) |

| | | | |
|---|---|---|---|
| ***CCL11/Eotaxin*** | 0.606788 | Inflammation | (Parajuli et al., 2015) (Lin et al., 2021) |
| **CCL17/TARC** | 0.797753 | Inflammation | (Achuthan et al., 2016) |
| **CCL20/MIP-3α** | 1.272791 | Inflammation, ERK inducer | (Zhang et al., 2017) |
| **CCL21/6Ckine** | 0.890857 | Inflammation | (Johnson & Jackson, 2010) (Damås et al., 2009) |
| **Clusterin** | 0.72376 | Inflammation, phagocytosis, ERK suppressor | (Weng et al., 2021) (Ungsudechachai et al., 2022) (Zhong et al., 2018) |
| **CNTF** | 1.23513 | Inflammation | (Hu et al., 2020) |
| **CXCL2/GROβ/MIP-2/CINC-3** | 1.243133 | Inflammation, phagocytosis, ERK inducer | (Boro & Balaji, 2017) (Lee et al., 2021) (Ha et al., 2010) |
| **Cyr61/CCN1** | 0.826287 | Anti inflammation | (Löbel et al., 2012) |
| **Cystatin C** | 1.34812 | Inflammation, ERK inducer | (Dutta et al., 2012) (Zhang et al., 2019) |
| **DPPIV/CD26** | 1.26338 | Inflammation, phagocytosis, ERK inducer | (Király et al., 2018; Ta et al., 2010) |
| **Galectin-3** | 1.379243 | Inflammation, phagocytosis | (Liu et al., 2022) (Sano et al., 2003) |
| **GM CSF** | 1.670754 | Inflammation, phagocytosis, ERK inducer | (Bozinovski et al., 2002) (Pinder et al., 2018) |
| **IGF 1** | 0.677557 | Anti-inflammatory, ERK inhibitor | (Sun et al., 2020) (Herrera et al., 2021) |
| **IGFBP3** | 1.51536 | ERK inducer | (Han et al., 2014) |
| **IGFBP6** | 1.311869 | Inflammation | (Chesik et al., 2004) |
| **IL-3** | 1.297589 | Inflammation | (Gebicke-Haerter et al., 1994) |
| **Jagged 1** | 1.31703614 | Inflammation, phagocytosis | (Wen et al., 2021) |
| **MMP2** | 1.376026 | Aβ degrading enzymes | (Dorandish et al., 2021) |
| **MMP9** | 1.335152 | Aβ degrading enzymes | (Dorandish et al., 2021) |
| **Neprilysin/CD10** | 1.442281397 | Aβ degrading enzymes | (Hafez et al., 2011) |
| **NOV/CCN3** | 1.47381373 | Inflammation | (Jia et al., 2017) |
| **OSTEOPONTIN** | 1.300466 | Inflammation | (Lalwani et al., 2023) |

| | | | |
|---|---|---|---|
| **PDGF** | 1.411737 | Inflammation | (Yang et al., 2016) |
| **RAGE** | 1.551495 | Phagocytosis receptor involved in inflammation | (Fang et al., 2010) |
| **Resistin** | 1.208339804 | Inflammation | (Azzam et al., 2020) |
| **RGM-A** | 1.664584095 | Inflammation | (Oda et al., 2021) |
| **SCF** | 1.658554449 | Inflammation, phagocytosis. ERK inducer | (Gao et al., 2022) (Wang et al., 2019) (Terashima et al., 2018) |
| **Serpin E1/PAI-1** | 1.370218952 | Inflammation, ERK inducer | (Pu et al., 2022) (Y. Wang et al., 2023) |
| **TIM-1/KIM-1/HAVCR** | 1.464351183 | Inflammation | (Zheng et al., 2019) |
| **VCAM-1** | 1.362232 | Inflammation, phagocytosis, ERK inducer | (Ishidome et al., 2017), (Kong et al., 2018) (Abdala-Valencia et al., 2011) |
| **WISP-1** | 1.506957 | Inflammation, ERK inducer | (Wang et al., 2022) (Chen et al., 2016) |

**Additional file 3:** Table showing fold change of differential secretion of proteins associated with microglial inflammation and phagocytic induction in response to Aβ as observed in cytokine array experiment

**Additional file 4**

| **TEST** | | **No of animals** | **Males** | **Females** |
|---|---|---|---|---|
| **Locomotion** | **WT** | **10** | **4** | **6** |
| | **5xFAD** | **9** | **4** | **5** |
| | **5xFAD+ rrICAM-1** | **9** | **4** | **5** |

| | | | | |
|---|---|---|---|---|
| | **5xFAD+ rrICAM-1+ lif** | **8** | **4** | **4** |
| **NOR** | **WT** | **8** | **4** | **4** |
| | **5xFAD** | **9** | **4** | **5** |
| | **5xFAD+ rrICAM-1** | **11** | **6** | **5** |
| | **5xFAD+ rrICAM-1+ lif** | **8** | **4** | **4** |
| **CDFC** | **WT** | **7** | **5** | **2** |
| | **5xFAD** | **7** | **4** | **3** |
| | **5xFAD+ rrICAM-1** | **5** | **3** | **2** |
| | **5xFAD+ rrICAM-1+ lif** | **6** | **3** | **3** |

**Additional file 4:** List of animal number and sex. NOR=novel object recognition, CDFC= cue dependent fear conditioning
